# Image processing techniques for high-resolution structure determination from badly ordered 2D crystals

**DOI:** 10.1101/113415

**Authors:** Nikhil Biyani, Sebastian Scherer, Ricardo D. Righetto, Julia Kowal, Mohamed Chami, Henning Stahlberg

## Abstract

2D electron crystallography can be used to study small membrane proteins in their native environment. Obtaining highly ordered 2D crystals is difficult and time-consuming. However, 2D crystals diffracting to only 10-12 Å can be prepared relatively conveniently in most cases. We have developed image-processing algorithms allowing to generate a high resolution 3D structure from cryo-electron crystallography images of badly ordered crystals. These include *movie-mode unbending*, refinement over sub-tiles of the images in order to locally refine the sample tilt geometry; implementation of different CTF correction schemes; and an iterative method to apply known constraints in the real and reciprocal space to approximate amplitudes and phases in the so-called missing cone regions. These algorithms applied to a dataset of the potassium channel MloK1 show significant resolution improvements to approximately 5Å.

**Abbreviations:**

2D: two dimensions / dimensional
3D: three dimensions / dimensional
*Amp*: amplitude
cAMP: cyclic adenosine monophosphate
CCD: charge coupled devices
CMOS: complementary metal-oxide-semiconductor
CNBD: cyclic nucleotide-binding domain
cryo-EM: cryo-electron microscopy
CTF: contrast transfer function
DED: direct electron detector
DQE: detector quantum efficiency
EM: electron microscope
*FOM*: figure-of-merit
*Pha*: phase
SNR: signal-to-noise ratio

## 1 Introduction

Direct electron detectors (DEDs) equipped with a radiation-hardened complementary metal-oxide-semiconductor (CMOS) sensor (Bammes et al., 2012; Glaeser et al., 2011a; Milazzo et al., 2011) tremendously increase the signal-to-noise ratio (SNR) of cryo-electron microscopy images. Instead of converting incoming electrons into light via a scintillator, these new devices detect them directly, which significantly reduces detector background noise. Beside the superior detector quantum efficiency (DQE) of DEDs (Ruskin et al., 2013), these detectors feature an enhanced sensor readout frequency making counting of single electrons possible to perform and resulting in recording of images with minimal dark background signal. This, along with the fast readout speed, allows recording image frames within exposure times that are short enough to prevent physical drift of the specimen stage. Consequently, image stacks, also called movies, of the sample can be recorded while it is continuously exposed to the electron beam. Algorithms like Zorro (McLeod et al., 2017), MotionCorr (Li et al., 2013b), MotionCor2 (Zheng et al., 2016), Unblur (Brilot et al., 2012), or alignparts_lmbfgs (Rubinstein and Brubaker, 2015) efficiently measure and correct for translational offsets between dose-fractionated movie-frames recorded on DEDs. The impact of sample movements such as those caused by specimen stage-drift, can thereby be strongly reduced, which increases the efficiency of data acquisition significantly. These new hardware and software developments have led to several recent breakthroughs in high-resolution cryo-EM 3D structure determinations that reach and surpass atomic resolution (Danev and Baumeister, 2016; Liu et al., 2016; Merk et al., 2016). Thus, cryo-EM has become a powerful and fast method for atomic-resolution protein structure determination.

2D electron crystallography is a branch of cryo-EM that is used to determine the electron density maps by imaging 2D crystals of desired biological structures (Stahlberg et al., 2015). Studying a protein system in form of 2D crystals is advantageous if (i) the hydrophobic core of a fragile membrane protein is to be contained in full lipid bilayer, thereby providing a native environment to the membrane proteins, (ii) the protein to study is too small for single particle cryo-EM studies, or (iii) the biological function of the protein is tied to a 2D crystalline arrangement. In such cases, the creation of homogenous sample with high crystallinity becomes important to generate a high-resolution structure by 2D electron crystallography. Unfortunately, obtaining highly ordered crystals is tedious and time consuming, since it involves testing protein stability in various detergents along with varying crystallization conditions. In several cases, however, 2D crystals diffracting to only 10-12 Å can be prepared relatively conveniently. "Bad" 2D crystals are easier to produce.

Besides sample quality, a beam-induced specimen movement further reduces the crystalline order, especially when the images of highly tilted 2D crystal samples are to be recorded (Glaeser et al., 2011a). The behavior of a biological sample under the electron beam was investigated in multiple studies (Bai et al., 2013; Brilot et al., 2012; Campbell et al., 2012; Glaeser et al., 2011a; Veesler et al., 2013). By analyzing large virus or ribosome particles movements during the exposure, these studies reported locally correlated movements that differ among the particles of an image. Such locally varying movement was also observed for 2D crystal samples before (Anchi Cheng, The Scripps Research Institute, La Jolla, CA, USA, personal communication). It is commonly believed that these local movements are due to irreversible deformations of the ice layer and also the sample, caused by the electron beam (Brilot et al., 2012). The amount of beam-induced motion depends on the microscope settings and the properties of the sample and grid. For 2D crystals,Glaeser *et al.* proposed firmly attaching the crystals to an appropriate strong and conductive support in order to overcome beam-induced sample movements (Glaeser et al., 2011a). However, such treatment may not be optimal for membrane proteins that show structural alterations when adsorbed to carbon film. A more general approach is to allow the 2D crystal membrane to move under the beam, and then computationally retrieve the structural data from recorded movies. Here, algorithms accounting for universal motion of all movie-frames, *e.g.,* caused by stage drift, cannot correct for spatially varying beam-induced motion as they neglect locally varying movements within the frames. An image processing approach that locally treats sub-regions of a 2D crystal sample in each movie frame is needed.

Another resolution-limiting factor is electron beam-induced radiation damage. A high cumulative electron dose destroys fine structural details within the proteins, whereas a low cumulative electron dose preserves the high-resolution information but reduces the SNR of the recorded image. Before detectors capable of movie-mode imaging were available, the full electron dose was captured within one single image and the EM operator had to carefully choose the electron dose depending on the target resolution. In 2010, before the advent of DEDs, Baker *et al.* recorded dose-fractionated image series of crystals and analyzed the resolution dependent vanishing of computed diffraction spots. Their findings suggest to record dose-fractionated image series of the sample and add these frames together using resolution-dependent frequency filters, in order to optimally exploit each image frequency at its maximal SNR (Baker et al., 2010), as is now routinely done in single particle cryo-EM image processing. The analysis by Baker et al., using crystalline material, resulted in a higher beam sensitivity, than the analysis of Grant & Grigorieff (Grant and Grigorieff, 2015), who used single particles for their study of optimal beam doses and related B-factor resolution filters. This might be caused by the fact that while single particles suffer within their protein structure from beam damage, a 2D crystal will in addition also suffer in crystal order due to beam damage. For this reason, a specific electron dose on a 2D crystal will lead to faster resolution loss in diffraction quality of recorded images, than the same dose would do to single protein particles. This resolution loss on 2D crystals, however, might be partly recoverable with appropriate image processing, if finer local movements of image segments can be traced.

The fact that a 2D crystal contains many regularly arranged identical copies of the same protein, allows electron crystallography to improve the SNR by using Fourier-filtering methods. Processing individual micrographs of 2D crystals generally involves the following six steps (Arheit et al., 2013a; Arheit et al., 2013b; Arheit et al., 2013c): (i) Defocus and (ii) tilt-geometry estimation, (iii) lattice determination, (iv) correcting for crystal imperfections, termed *unbending*, (v) Fourier extraction of amplitude and phase values from computed reflections, (vi) Contrast transfer function (CTF) correction and (vii) generating a projection map of one unit cell at higher SNR. If this procedure is applied to the drift-corrected average gained from a movie-mode exposure of a 2D crystal, it does not account for locally varying beam-induced sample movements. Additionally, the CTF determination and correction is done after the unbending procedure. With the images obtained from new detectors, Thon rings up to 3 Å can be correctly identified. GCTF (Zhang, 2016) or CTFFIND4 (Rohou and Grigorieff, 2015) can quickly and precisely measure the CTF and defocus of a recorded image respectively movie. This creates a possibility to correct the CTF as a preprocessing step before the unbending procedure.

Once the projection maps for all images are generated, they are merged in Fourier space using the *Central Projection Theorem*, and the target 3D density profile (3D map) is reconstructed by inverse Fourier transform. The extent to which a sample can be tilted in the electron beam is limited to about 60 degrees. This is because the thickness of the irradiated layer increases with the tilt angle, decreasing the transmitted electron beam so that there is hardly any signal at higher tilt angles. Other factors such as sample movements and varying defocus also complicate the situation. As a consequence, slices with tilt angles beyond this limit, are missing from the reconstructed 3D Fourier space. Together, these missing slices correspond to a conical volume known as the “missing cone”. In real space, the missing cone means that reconstructed densities are smeared out in the vertical (Z-) direction (**Supplementary Fig. 1**). Data can also be missing in other Fourier space regions, depending on the tilt sampling. An infinite number of sample tilts would be required to completely fill the 3D space.

In this study we developed image-processing algorithms and techniques allowing to generate a high resolution 3D structure from cryo-EM images of badly ordered 2D crystals. These include: (i) *movie-mode unbending* that corrects for locally varying beam-induced sample deformations and computationally applies electron dose-dependent resolution filters, (ii) *tiled image processing*, by dividing images into sub-images called tiles, minimizing the errors that arise when using of same tilt angle for the whole image, (iii) application of different choices of CTF correction schemes for 2D-electron crystallography with the aim of increasing resolution, and (iv) *shrinkwrap refinement*, to fill in missing amplitudes and phases in the 3D volume. The new procedures are implemented as additional scripts in Focus (Biyani et al., 2017), previously 2dx (Gipson et al., 2007b), and are available for download at http://www.2dx.org or http://www.focus-em.org. We present the application of the new techniques to an experimental dose-fractionated membrane protein dataset recorded on a FEI Titan Krios equipped with a Gatan K2 summit detector, which yielded an improved 3D reconstruction of the potassium channel MloK1.

## 2 Theory

The field of electron crystallography of membrane proteins was created through the work of Richard Henderson and Nigel Unwin (Henderson and Unwin, 1975; Unwin and Henderson, 1975). Their developed algorithms were made available to the public in the so-called MRC programs for image processing (Amos et al., 1982; Crowther et al., 1996b; Henderson et al., 1986; Henderson et al., 1990), which were streamlined and automated by the 2dx software package (Arheit et al., 2013a; Arheit et al., 2013b; Arheit et al., 2013c; Gipson et al., 2007a; Gipson et al., 2007b). The MRC software approach was thereby based on the idea that the sample is a set of several perfectly ordered and flat 2D crystals, which are imaged in a cryo-EM instrument under different tilt angles. Such images are then computationally unbent for crystal lattice distortions, followed by transformation into Fourier space, evaluation of amplitudes, phases, and background values for each crystal reflection, and these data are then merged into a 3D Fourier volume, following the "central section theorem", which states that the projection of a sample imaged at a certain tilt angle corresponds to a central slice in 3D Fourier space under the same tilt angle. This Fourier space central section approach assumes that an entire image of a 2D crystal shows each fraction of the protein crystal under the identical tilt geometry. This assumption is voided, if the 2D crystalline sample was not on a flat plane (as is normally the case when the crystalline membrane is freely suspended in a vitrified buffer layer without underlying carbon film support). Or, even if the crystal were on a perfectly planar surface, any in-plane distortions in the 2D crystalline arrangement will lead to varying tilt geometries for different unit cells in the same crystal, if the sample were imaged under higher sample tilt. This fact leads to a resolution limitation, when badly ordered 2D crystals are to be imaged under higher tilt.

### 2.1 Tilt geometry definition in 2D electron crystallography

The tilt geometry of a 2D crystal is in electron crystallography parameterized with the variables TLTAXIS and TLTANG. The MRC image convention has the first point of the image in the lower left corner indexed as [0,0], with the x-axis pointing right and the y-axis pointing up. TLTAXIS defines the orientation of the sample tilt axis, measured from the horizontal X-axis (pointing right in the image) to the tilt axis with positive values meaning counter-clockwise. TLTAXIS is defined as between −89.99999° (pointing almost down) and +90.0° (pointing straight upwards). TLTANG defines the tilt angle of the specimen, and can take any value between −89.999 and +89.999 degrees. Due to the physical limitations affecting a planar 2D crystal in an electron microscope, a tilt angle of 90° is not possible. If the TLTAXIS is horizontal and underfocus in the image gets stronger further up in the image (*i.e.,* at locations with higher y coordinates), then TLTANG is positive.

In electron crystallography, the tilt geometry from the point of view of the 2D crystal is defined by the parameters TAXA and TANGL. TAXA is the relative rotation of the unit cell of the given 2D crystal in 3D Fourier space, and TANGL is equal to TLTANG, but its sign alters depending on the orientation of the crystal on the tilted specimen plane, or the handedness of the assigned lattice vectors (Arheit et al., 2013a). These parameters TAXA and TANGL are only relevant for 3D merging in electron crystallography, and are not further discussed here.

Other software packages such as SPIDER (Shaikh et al., 2008), FREALIGN (Grigorieff, 2016), EMAN2 (Bell et al., 2016) or RELION (Scheres, 2012) define the tilt geometry differently. The relationship between the electron crystallography tilt geometry adopted in the MRC and 2dx software packages, and the SPIDER convention of Euler angles adopted in many single particle reconstruction packages, is the following:

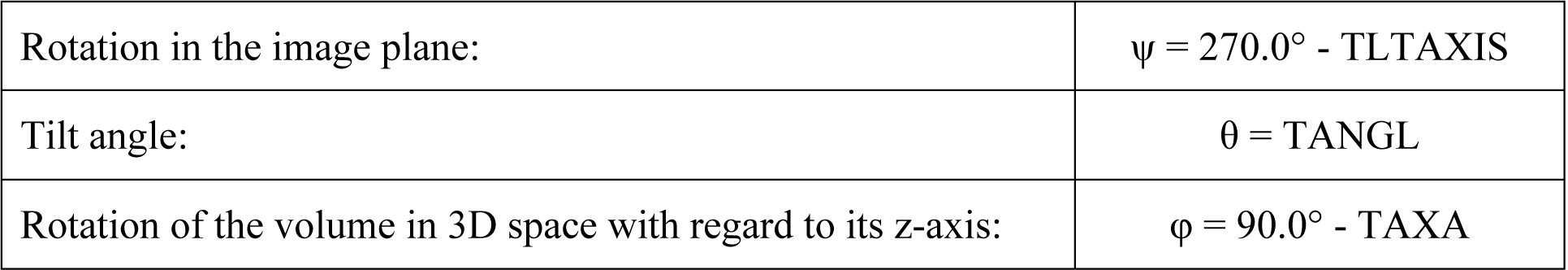

|  |  |
| --- | --- |
| Rotation in the image plane: | $\psi = 270.0^\circ - \text{TLTAXIS}$ |
| Tilt angle: | $\theta = \text{TANGL}$ |
| Rotation of the volume in 3D space with regard to its z-axis: | $\phi = 90.0^\circ - \text{TAXA}$ |

### 2.2 Tiled image processing

To minimize these effects, we propose to subdivide the image into smaller parts called tiles. The subdivision, ideally, should target a tile size that is small enough to identify the correct local tilt angles throughout the image. But a smaller tile will have less signal to noise ratio, which will make correct 3D merging of the evaluated amplitude and phase data more challenging. A suitable tile size such as 1024×1024 pixels or larger should therefore be chosen such that there is enough signal in the image. To reduce the Fourier artifacts that arise due to the edges, cropped tiles are edge-tapered, so that some image segments are lost. To reduce data loss, tiles are therefore extracted with overlapping edges. Once the image is properly divided into tiles (e.g., 5 tiles in one dimension, resulting to 25 tiles), each of these tiles is considered an individual image (**Figure 1**). Most parameters for the tile images are inherited from the parent image, while the local defocus is calculated based on the specific height of the central pixel in the newly cropped tile. The parameters for each tile are then refined in iterative cycles, whereby the tilt geometry, defocus and astigmatism, and also beam tilt can be allowed to vary within constraints from the given starting values. This process increases in the above example of 5×5 tiles the number of 2D crystal datasets by 25, so that a much larger number of reflections has to be submitted to the 3D refinement and 3D merging steps (**Figure 1**). This approach is computationally demanding; to increase the efficiency, recalculation of parameters such as defocus can be search limited to the a value corrected for the predicted Z-height of the tile, while limiting the search range to a more narrow range.

**Figure 1:**
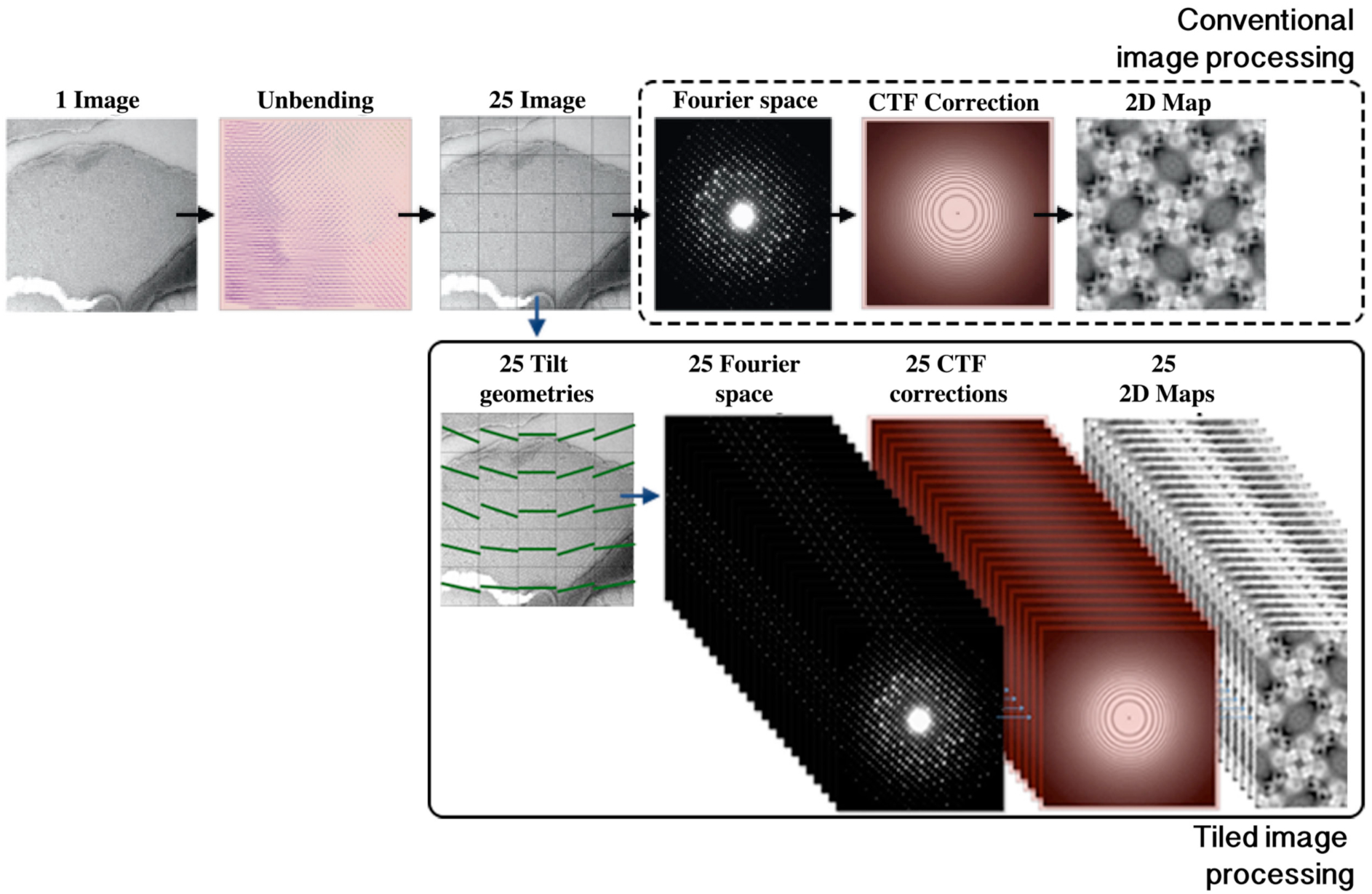
Tiled image-processing scheme. When crystals are not flat, the parameters like tilt-geometries and defocus vary along the image. To reduce this problem, images are divided into various overlapping tiles and the tiles are individually processed to get many 2D Maps instead of one.

### 2.3 CTF correction strategy

Correction of the Contrast Transfer Function (CTF) of the electron microscope is necessary in order to restore structural information, especially at higher resolutions, that would otherwise be lost during the image processing and merging steps. In the absence of beam-tilt and in case of parallel illumination, the electron microscopy projection of the biological structure in the image is convoluted in real-space with the point-spread function (PSF) of the microscope, which is equivalent to a multiplication of the Fourier transformation of the image with a real-valued CTF function. In case of beam-tilt or non-parallel illumination, the PSF in real space has resolution-dependent delocalization features, or the CTF in Fourier-space becomes a complex-valued function (Glaeser et al., 2011b). The effects of this are not further discussed here.

The real-valued CTF shows oscillatory rings, called Thon rings (Thon, 1966), which refer to the maxima of the CTF, which are interrupted by zero-crossings of the CTF. Their diameter depends on the applied defocus under which the image was recorded. In case of a tilted sample, that defocus varies across the image. A reliable measurement of the tilt geometry parameterized by the variables TLTANG and TLTAXIS, and also of the magnification is therefore mandatory to be able to fully correct for the effect of the varying defocus.

The tilt geometry is in a first step determined by a geometric evaluation of the defocus gradient, as described in detail in (Arheit et al., 2013a). The tilt geometry is refined in a second step, based on the distortions of the measured 2D crystal lattice, and its deviation from the non-tilted 2D crystal lattice. This will only give reliable measurements for specimens at higher tilts (*i.e.,* TLTANG > 25°). In a third step, the tilt geometry is refined during merging, by comparing the amplitude and phase data from each image with the current 3D reconstruction. This refinement of the tilt geometry can be separately done also for the sub-tiles of the images, as discussed in the previous section.

All these tilt geometry refinement steps are unable to detect or eliminate the consequences of a wrong pixel size. The defocus gradient measurement takes the current definition of the pixel size into account. If for example that pixel size was quantified 1% too small, then the defocus measurement in the corners of the image will have minimally different values, but if assigned to locations too far apart from each other, results in an underestimation of the tilt angle (TLTANG). As a result, the determined 3D reconstruction would be too short in the z direction, while being too wide by the same ratio in the x and y directions. This problem of resulting anisotropic magnification does not apply to single particle cryo-EM, or to helical image processing, where a magnification error would simply result in a wrong scale of the final reconstruction.

Anisotropic scaling of a final map in 2D electron crystallography can be recognized and corrected, if sufficient resolution is reached to fit the pitch of alpha-helical structures in a final map to expected biological values. This, however, requires a final 3D resolution of better than 4.5 Å. At lower resolution, the aspect-ratio of a 3D reconstruction from electron crystallography therefore has to interpreted with care.

Correction of the effect of the CTF can be done in various ways in electron crystallography.

#### 2.3.1 CTF correction methods

CTF correction can be done by simple phase-flipping (phases of Fourier pixels on odd-numbered Thon rings are incremented by 180°, so that the phases are corrected but the amplitude oscillations in the original data remain), or by CTF multiplication (the Fourier pixels are multiplied with the CTF pattern ranging from −1 to +1, so that the phases are corrected but the amplitude oscillations in the original data are now even more exaggerated), or by Wiener filtration (the Fourier pixels are multiplied by CTF / (CTF^2^ + N^2^), so that the phases are corrected, and the amplitudes are partly re-established to their correct values. Here, *N* is a noise term that depends on the SNR of that image segment. A value of *N*=0.3 gives conservative results in practice.).

Richard Henderson et al. introduced the correction of the "tilted transfer function" (TTF), by first unbending the 2D crystal images, then Fourier transforming the unbent image, and convoluting the Fourier transform for each lattice spot with a pattern that represents the expected split spot profile at that location (Henderson et al., 1986). Only after convolution, the amplitude and background values are evaluated. This so-called TTF correction effectively eliminates all effects of the defocus gradient across the image, but assumes that the image has already been unbent and the structural information is contained in the precisely located Fourier reflections. Effects of beam tilt have to be addressed during 3D merging later. Effects of non-parallel illumination (*i.e.,* coma) cannot be corrected with this method. Effects of a varying tilt geometry throughout the image (*e.g.,* from a bent 2D crystal plane) also cannot be addressed with this method.

Ansgar Philippsen *et al.* developed a tilted contrast image function (TCIF), which is an extension of the TTF approach above (Philippsen et al., 2007). Philippsen et al. thereby showed that the transfer function of an electron microscope on tilted samples is only in a first approximation a convolution with a (varying) point spread function across the image. The complexity of the developed mathematics so far precluded the implementation of a computationally efficient reversion of the effects of the TCIF on cryo-EM images.

#### 2.3.2 CTF correction in stripes before unbending

We have developed software to correct the effect of the varying CTF in the real-space image before unbending. This is done by correcting the image with a static CTF profile by multiplication in Fourier space, and extracting only a narrow stripe in the direction of the tilt axis at the image regions, where that CTF agrees with the local defocus. The stripe width is thereby chosen so that the defocus error on the edges of the stripe does not result in a resolution limitation, following the estimations in (Zhang and Zhou, 2011). For example, if a target resolution of 3Å is to be reached, defocus tolerance would be less than 20 nm, which then defines the width and the number of stripes required in dependence of the estimated tilt angle. Assembling various such stripes eventually produces the fully, locally CTF corrected output image. Any effect of beam tilt is not considered, and would have to be addressed during 3D merging.

#### 2.3.3 CTF correction before vs. after unbending

CTF correction can be done before 2D crystal unbending, or after unbending, or in a combination of both. CTF correction before unbending has the advantage that delocalized features from extended point spread functions (PSF) in the real-space image are moved to their correct locations before the crystal unbending moves image segments around. If unbending is instead done first, followed by CTF correction later, then the unbending step risks to cut the ripples of the PSF, thereby loosing high-resolution data. For badly-ordered 2D crystals, this problem can become resolution limiting. CTF correction before unbending by Wiener filtration, on the other hand, reduces the SNR in the image, so that the unbending step cannot as reliably determine the crystal order defects.

#### 2.3.4 Hybrid CTF correction

We also have tentatively implemented a so-called "Hybrid CTF correction" algorithm, where the original image is corrected in two different versions in stripes for the varying CTF across the image: One copy of the image is corrected by CTF multiplication, and a second copy by CTF Wiener filtration. The first version then has a maximized SNR of the biological structures, and is ideal for determination of the crystal defects and the unbending profile. Attempts were made to measure the crystal distortions on the CTF-multiplied image, but then applying the unbending procedure to the Wiener-filtered image instead. In practice, however, the unbending profile determined on the CTF-multiplied copy of the image, was not found precise enough to unbend the Wiener-filtered image to high resolution. Better results were obtained by applying the unbending algorithm to the same image for which the unbending profile had been measured. As a consequence, CTF correction by simple phase flipping on stripes before unbending is generally recommended.

### 2.4 Movie-mode unbending

2D crystal images show different local movements within regions of a single 2D crystal during imaging (**Figure 2**). When recording movies with a total dose of 40 electrons / Å^2^, we observed local relative movements of up to 4 nm within micrometer-sized 2D crystal regions. This observation underlines the need for per-frame movie-mode image processing for high-resolution electron crystallography.

**Figure 2:**
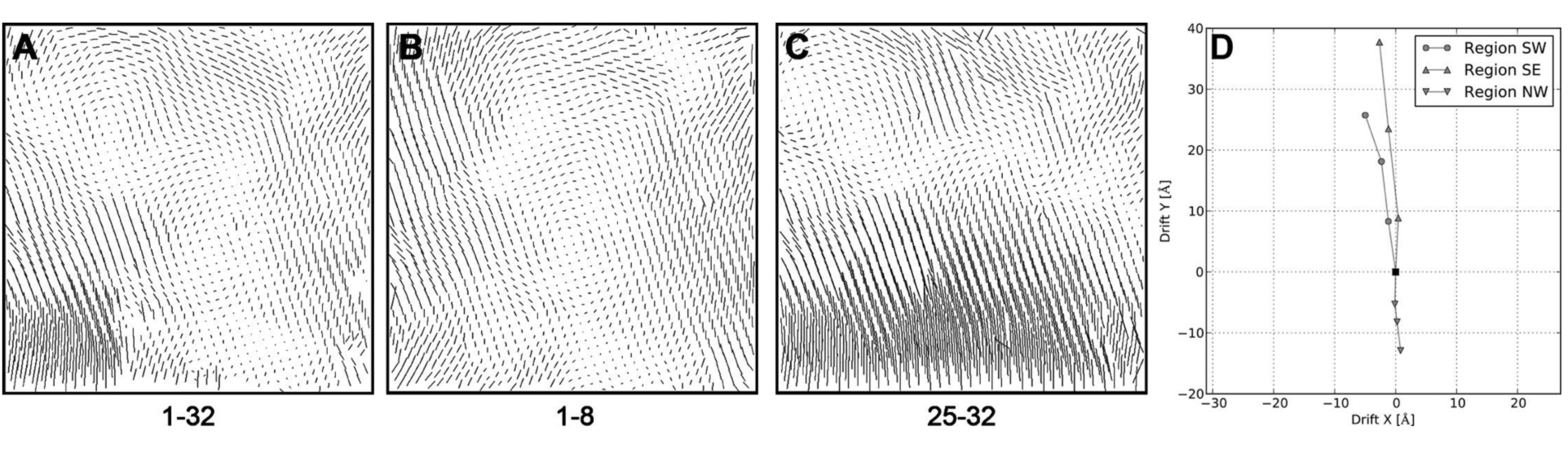
Electron dose dependent variation of the distortion-vector ERROR-fields showing the necessity of beam-induced motion-correction. (**A**) Unbending profile (4k × 4k pixels) obtained by processing the average image generated by real-space averaging of 32 movie-frames (out of the recorded 40 frames, after discarding the first two and the last 6 frames). For display purposes, the motion vectors have been elongated ten times. By grouping and average frames into super-frames, we obtained a new movie consisting of only four super-frames, showing the crystal at different exposure times (averaging frames 1-8, 9-16, 17-24, and 25-32, after discarding the first two and the last 6 frames). These four super-frames were then independently unbent with the algorithm described as MovieB. The distortion-vector ERROR field of the first average (frames 1-8) is shown in (**B**), the one for the fourth average (frames 25-32) in (**C**). (**D**) Drift-profiles quantifying the averaged local movements over the whole sequence of super-frames relative to the first frame average. Fifty neighboring trajectories were averaged for each quadrant of the images. The drift of the northeast quadrant of the crystal was omitted because of visualization difficulties due to negligible local drift (< 2 Å). The lower area (southern quadrants) of the crystal is drifting upwards during the exposure, whereas the movements of the upper part are smaller and in another direction. An approach that corrects for sample movements at frame level by translationally aligning entire frames is not able to correct for such inhomogeneous and electron dose-dependent local crystal deformations.

The fundamental idea of movie-mode unbending is to process each frame of a movie individually and merge the data from the unbent frames later. This requires (i) correctly exploiting the smooth motion of proteins in consecutive frames, (ii) accounting for radiation damage during the increasing exposure, (iii) dealing with the extremely low SNR of one movie-frame, and finally (iv) preventing overfitting.

The classical unbending procedure is briefly described here followed by the description of two alternative movie-mode unbending algorithms, as schematically detailed in **Figure 3**. The algorithm starts with the conventional unbending of the whole-frame drift-corrected and averaged single image, using the conventional Unbend2 algorithm. During this step, an unbending reference (termed "Ref." in **Figure 3A**) and also an unbending profile are determined.

**Figure 3:**
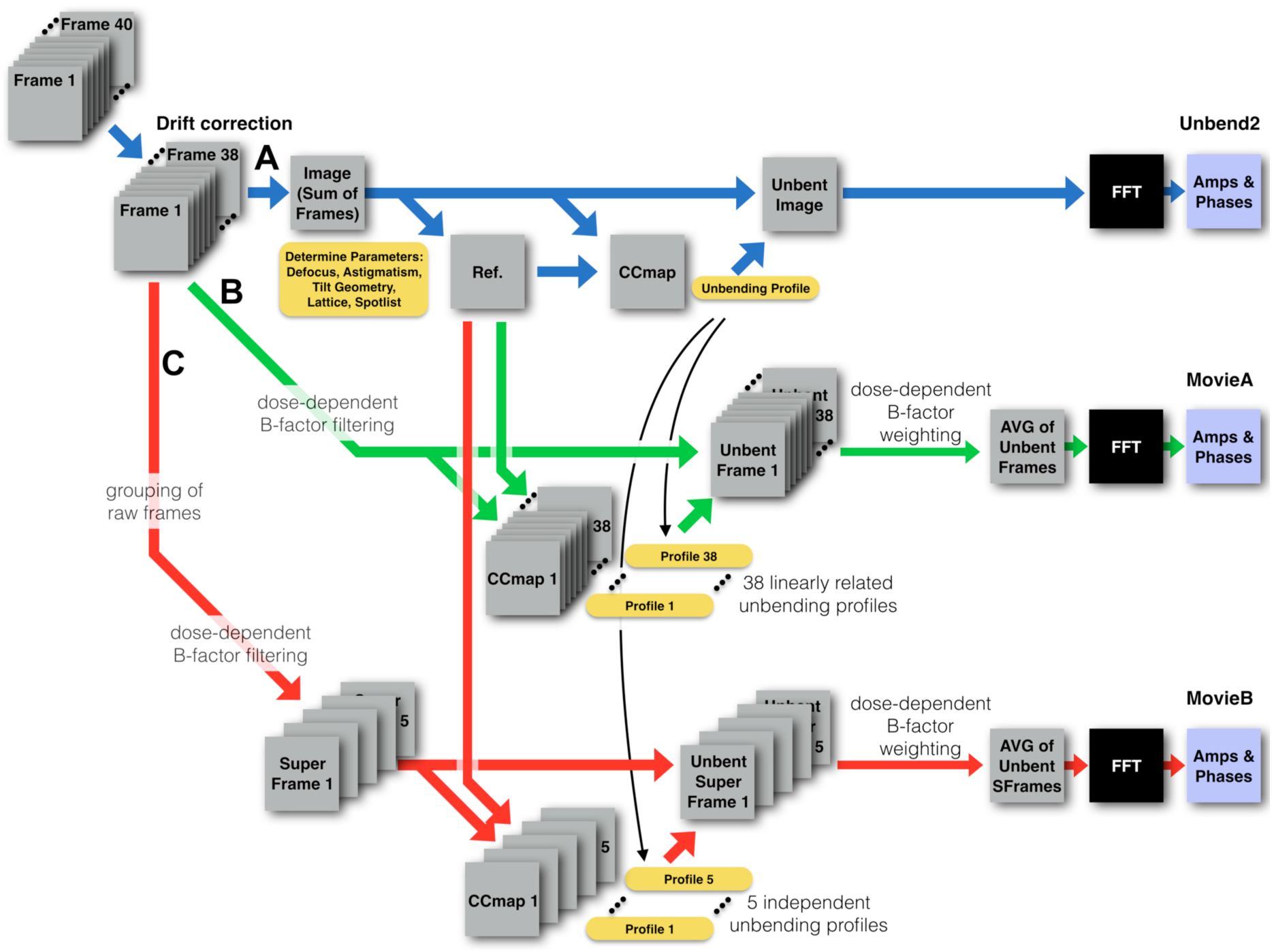
Movie-mode unbending algorithms. The recorded image stack of 40 frames is whole-frame drift corrected, *e.g.,* using Zorro (McLeod et al., 2017) or MotionCorr2.1 (Li et al., 2013b). Here, the first two frames are typically discarded due to elevated drift. (**A**) The remaining 38 frames are averaged and the single, resulting image is conventionally processed by unbending, resulting in the dataset termed "Unbend2" result. (**B**) A movie-mode processing algorithm termed MovieA is then applied to the whole-frame drift-corrected stack. MovieA unbends each of the 38 frames individually, using as unbending reference the reference from the Unbend2 branch (labeled “Ref.”). In order to be less sensitive to noise, the 38 unbending profiles for the 38 movie frames are established by assuming that each 2D crystal unit cell can only drift along a linear trajectory of constant speed during the entire movie. This constraint allows unbending each of the 38 frames individually, resulting in the final dataset termed “MovieA”. (**C**) An alternative movie-mode processing algorithm termed MovieB is also applied to the remaining 38 movie frames. This approach consists of grouping the frames into so-called super-frames, e.g., by combining groups of 7 original frames into one super-frame, resulting in 5 super-frames. These then provide sufficient signal-to-noise ratio to allow the determination of individual unbending profiles for each super-frame. The 5 super-frames are then unbent, and the data combined, resulting in the final dataset termed “MovieB”. Both, MovieA and MovieB “learn” from the unbending profile in the Unbend2 run. Also, both movie-algorithms apply electron-dose dependent B-factor filtering of high-resolution terms in each frame, the cumulative decay of high resolution terms is later corrected during combination into a merged dataset for this movie.

#### 2.4.1 Classical unbending procedure

The central step in the processing of one single image of a 2D crystal is correcting crystal imperfections, termed unbending (Arheit et al., 2013a). A Fourier-filtered reference is generated based on the crystallographic lattice determined in reciprocal space, and cross-correlated with the micrograph. Peaks in the resulting cross-correlation profile (localized by the MRC program Quadserch (Crowther et al., 1996a)) correspond to crystallographic unitcell locations. The MRC program CCUnbend is subsequently used to translationally adjust small patches (e.g., 25×25 pixels in size) of the micrograph to maximize the regularity of the crystal in order to achieve high-resolution projection maps. The image unbending is usually refined over several rounds, resulting in an improvement and sharpening of the computed diffraction spots. To avoid malicious noise accumulation or the domination of reference bias, the raw image is thereby never unbent multiple times, but instead a so-called distortion-vector field or “ERROR field” is continuously improved. The ERROR field obtained in the previous round of unbending is used as the starting solution for the current round of refinement. Generally, after three rounds of refinement, the final ERROR field is applied to unbend the raw micrograph in one step. The Fourier-transform of the unbent image is then evaluated to obtain the amplitude and phase values at the lattice reflection sites, and the data are CTF-corrected.

#### 2.4.2 Algorithm: MovieA

We implemented a movie-processing algorithm termed "MovieA", which is attempting to unbend each movie-frame individually (**Figure 3B**). This, however, generally fails for image frames that have very low electron counts (in our settings < 2 e^−^/pixel), because the remaining signal-to-noise ratio of individual frames no longer reliably allows the determination of the unbending profile. Instead, we implemented an algorithm that starts from the unbending profile of the averaged image (Unbend2 profile), and searches each crystal lattice node for a linear trajectory, along which these would have drifted with constant speed during the recording of the frames. The obtained set of drift vectors is further compared with the drift vectors of neighboring crystal unit cells: if one drift-vector significantly deviates from that of its closest neighbors, this vector and its corresponding image patch are deleted from the dataset. Deletion of such an image patch is done by replacing the corresponding image location in all frames with a Gaussian blob of average grey.

This linear-movement constraint assumes that the movement of every crystal lattice node is only linear and only in one specific direction. Any crystal part that moves first in one direction, and then turns to move into a different direction, cannot be correctly fitted with this approach.

The determined unbending profiles are then used to unbend each movie frame individually, and the resulting unbent frames are averaged in real space, Fourier-transformed, and evaluated for their amplitudes and phases. To account for electron-beam induced resolution loss, individual frames are B-factor filtered before unbending, and the final, unbent and averaged map is corrected for the cumulative B-factor filtration profile before evaluation of Amplitudes and Phases, as also done in RELION (Scheres, 2012), or MotionCor2 (Zheng et al., 2016), or Unblur (Brilot et al., 2012), and further detailed below.

#### 2.4.3 Algorithm: MovieB

We also implemented a second, alternative unbending algorithm, termed "MovieB" (**Figure 3C**). In this algorithm, the electron-dose dependent B-factor filtered movie frames are grouped into fewer so-called super-frames, by, e.g., grouping each 7 raw frames into one super-frame. The resulting movie is composed of much fewer super-frames, each of which now has an improved signal-to-noise ratio. These super-frames can then be individually unbent, using the high-contrast “Ref.” from Unbend2, and as starting template the unbending profile from the Unbend2 run. The resulting, individually unbent super-frames, are then combined in Fourier space, corrected for the B-factor filtration, and evaluated, resulting in a set of amplitudes and phases termed MovieB.

In the above described algorithms for MovieA (linearly correlated unbending profiles) or for MovieB (higher-contrasted super-frames), several parameters such as defocus, tilt geometry, crystal lattice, as well as the reference map for cross correlation with each movie frame, are taken from the first processing run termed Unbend2. The program *2dx_quadserch* was extended to be able to iteratively refine the determined ERROR field. When processing the first frame of a movie stack, we use the ERROR-field generated by the script *“Unbend II”* as the starting ERROR field for refinement. For all subsequent frames, the ERROR field generated while processing frame (n-1) is used as the starting ERROR field for the peak localization when unbending frame (n). This approach exploits the smooth correlation between the crystal distortions among subsequent movie-frames to ensure a continuous and physically comprehensible motion-correction. Finally, the MRC program *CCUnbend* is used to correct each movie frame according to its corresponding ERROR field by shifting small image patches. The unbent images are then averaged, yielding one combined output image. Using the same global reference image for all frames ensures that the unbent frames from UnbendII, MovieA, and MovieB runs are in register, which is helpful for later 3D merging. The MRC program *MMBoxA* is finally employed to evaluate the averaged unbent frames for all three algorithms, and to produce one APH-file each from the averaged frames. Among other things, these APH-files contain amplitude, phase, and SNR information for all computed diffraction spots. After correcting for CTF-effects (if not done before), a 2D projection map is generated for visual inspection, and finally the produced APH-files can be used for 3D merging.

#### 2.4.4 Accounting for resolution dependent radiation damage

As demonstrated in detail for catalase crystals by Baker et al. (Baker et al., 2010), electron beam damage is a crucial limit to high-resolution in cryo-EM. Based on the electron dose-dependent fading of diffraction spots, they showed how the optimal electron dose depends on the targeted resolution. For instance, to record data with an optimal SNR, a target resolution of 3 Å requires a cumulative electron dose of 11 electrons per Å^2^, while a target resolution of 27 Å requires a total dose of 22 electrons per Å^2^. Generally, a higher cumulative electron dose destroys the high-resolution details, but amplifies low-resolution features, such as overall shape and position of the particles.

DED movie-mode data acquisition allows dose fractionation, *i.e.,* the distribution of a high electron dose over a large number of movie-frames. In this work, we recorded data with a cumulative dose of 40 electrons/Å^2^ distributed over 40 movie-frames. Averaging a certain number of movie-frames during image processing allows average images corresponding to various desired electron doses to be generated. This has for example made it possible to assess the impact of electron beam damage on different amino acids (Allegretti et al., 2014; Campbell et al., 2012). Nevertheless, such *a-posteriori* averaging of movie-frames still suffers from the trade-off between the optimal electron dose required to obtain the low and high-resolution information present in the images. Investigations of the dose-dependent fading of computed diffraction spots of thin 3D catalase crystals with *2dx* were done in various studies (Baker et al., 2010; Bammes et al., 2010). We subjected this test data set to the same measurements of dose-effects for lipid membrane 2D crystals (**Supplementary Fig. 2**), and found comparable dose-dependent resolution fading for 2D crystals as previously published for thin 3D crystals. However, as also discussed in (Grant and Grigorieff, 2015), the loss in computed diffraction must be caused by both, structural damage to the membrane proteins, and loss of crystal order due to local movement or rotation of unit cells. Assuming that the here presented crystal unbending and tile processing algorithms can partly compensate for the loss in crystal order, the resolution decay values determined for single particles in (Grant and Grigorieff, 2015) should more accurately describe the optimal dose-dependent resolution weighting for 2D crystals that are processed with movie-mode unbending. This assures that all structure frequencies contribute to the resulting projection map at their optimal, dose-dependent SNR.

After having omitted the first two frames from the recorded movies (*i.e.,* skipping the first 2 e^−^/Å^2^), the B-factor filters allow for the first few frames all spatial frequencies to fully contribute, whereas for later frames primarily lower frequencies contribute to the average. Following (Grant and Grigorieff, 2015), the exposure-dependent resolution filter for a movie with *n* frames will compute for each Fourier voxel *F(k)* a weighted Fourier voxel *F^W^(k)* by

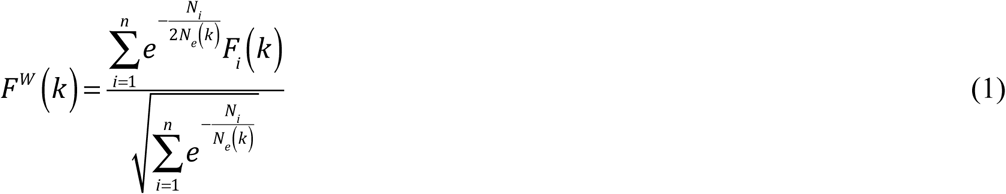

with *k* being the spatial frequency, *n* being the number of frames in the movie, *i* being an index identifying each frame, and *N_i_* being the cumulative numbers of electrons per square Å for frame *i*, and *N_e_* being the resolution-dependent critical electron exposure, and *F_i_(k)* being the Fourier component at resolution *k* in frame *i*.

The critical exposure term *N_e_(k)* in electrons per square-Angstroms is thereby calculated as

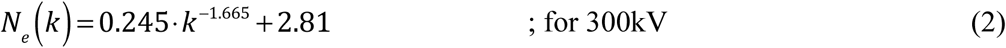

while for 200kV the critical exposure is estimated 25% lower, or

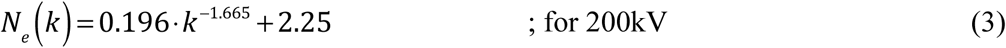

with *k* being the spatial frequency in 1/Å.

Application of the above equations allows dose-fractionated movies of 2D crystals to be processed with optimal weighting of frequency components.

### 2.5 Prevention of overfitting

#### 2.5.1 When grouping frames into super-frames

The tendency to mistake noise for interpretable structural elements can be major problem of iterative algorithms used in cryo-EM. This so-called *overfitting* arises from the combination of the low SNR generally present in cryo-EM images and over optimistic resolution estimations for the reference used as template in different stages of the data processing. When we applied the above algorithm to all movie-frames of an image-stack consecutively (electron dose of ~1 electron per Å^2^ per frame or lower) we observed a serious overfitting risk indicated by discontinuous distortion-vector-fields and artificial ripples in the resulting projection maps (**Figure 4**). Due to the low signal present in each frame, the cross correlation profile lacked distinct correlation peaks, which significantly reduced the reliability of the MRC program *Quadserch* to track the correct crystal unit cell locations. Increasing the ambiguity of cross correlation peaks can only be achieved by increasing the SNR of the raw data or the reference image. Using a larger reference would give a higher SNR for the cross-correlation map, but reduce the possibility to track fine crystal distortions, *i.e.,* any distortions that are smaller than the reference dimensions. On the other hand, increasing the SNR of the frames by using a higher electron dose per frame leads to beam-induced blurring within the frames caused by sample instabilities in the EM. Thus, a compromise has to be made. Here, in the algorithm termed "**MovieB**", we chose to average a given number of consecutive movie frames and process the resulting sub-frame averages. Such frame averages overcome the SNR-induced limitations described above. However, averaging multiple sub-frames and thus reducing the *“electron dose fractionation*” limits the possibility to capture all beam-induced motion. Nevertheless, the situation can be improved by applying a global frame-drift correction to each low SNR frame prior to the averaging step, in order to compensate for global stage drift and homogeneous beam-induced sample movement.

**Figure 4:**
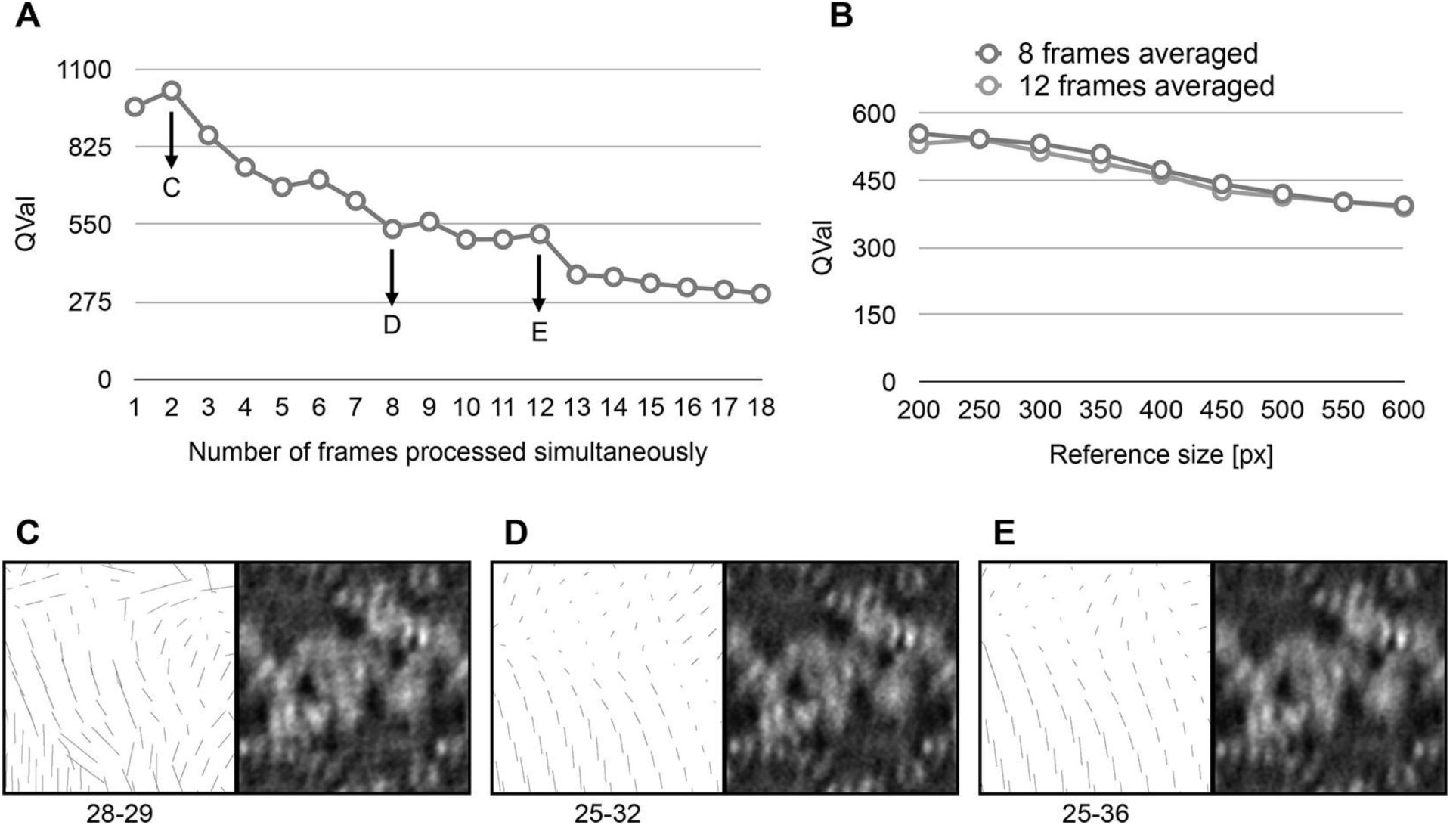
Development of movie-frame unbending for the MovieB algorithm, using the MloK1 test dataset. For this test dataset, classical processing resulted in a *QVal* of 232.2. (**A**) Relationship between the number of averaged movie-frames and the measured *QVal.* During drift-correction at frame level, the two first frames were removed and thus omitted from the movie-mode unbending. Three different regimes were found: *overfitting regime* when 1-7 frames were averaged, indicated by artificially high *QVals; working range* of the processing procedure when 8-12 frames were averaged; and *stiffness region* when 13-18 sub-frames were averaged, where the detection of beam-induced motion-correction is limited due to large number of averaged frames. (**B**) Impact of the reference size on the measured *QVal* analyzed for two different averaging schemes. (**C-E**) Visual comparison of the distortion-vector ERROR field (left) and the resulting projection map (right) when processing averages consisting of 2 frames (**C**), 8 frames (**D**), or 12 frames (**E**) were processed. The chaotic ERROR field in C is accompanied by striation artifacts in the image. D shows an optimal approach. E shows increased blurring in the image, due to a too smooth ERROR field.

This poses the question of how many frames or what electron dose per Å^2^ should optimally be averaged and processed together. We tested the movie-mode unbending algorithm with all conceivable numbers of averaged sub-frames and using the calculated parameter *QVal* (Gipson et al., 2007b) to assess the obtained quality (**Figure 4A**); a higher *QVal* value indicates that the image has more and/or sharper diffraction spots. In addition, we inspected the distortion ERROR fields and the projection maps obtained to see if overfitting could be visually detected (**Figure 4C-E**). Each of the movies analyzed consisted of 40 frames with a cumulative electron dose of 40 electrons/Å^2^. As cryo-EM samples have the tendency to move most at the beginning of the exposure, we completely omitted the first two frames. Furthermore, the last six frames were only included in the initial high-SNR average calculated after global drift correction of the individual frames. To avoid the effects of beam-induced sample damage, these six frames were not otherwise used. Hence, we fully processed 38 frames for MovieA, and only 32 frames for MovieB, covering a cumulative electron dose of ~32 electrons/Å^2^. Note that ensuring equal divisibility of the data for the analysis means that the total number of frames included depends on the number of frames grouped into super-frames. On analyzing the effect of averaging different numbers of frames into super-frames before movie-mode unbending, we observed three different regimes where the performance and reliability of the MovieB procedure differed significantly (**Figure 4A**). There were signs of overfitting when only a small number (1 to 7) of frames were averaged and processed, and the procedure was no longer able to resolve the beam-induced motion accurately enough if too many (13 to 18) frames were averaged. Averaging 8 to 12 frames gave optimal results. Movie-mode unbending of the resulting super-frame movies comprised of 3 or 4 averaged frames did not produce any signs of overfitting and still allowed beam-induced crystal alterations to be reduced. Although our observations were consistent over the whole MloK1 test dataset, the optimal batch size presumably depends on the sample as well as on the imaging conditions (structural contrast in the biological structure, acceleration voltage and applied defocus of the TEM, pixel size and type of utilized detector). It must be stressed that although the final movie used for movie-mode unbending was only comprised of 4 high SNR frame averages, the 40 individual frames originally recorded were essential to allow global drift-corrections to be made over a fine dose raster. This important correction would not be possible if just 4 higher dose, and thus higher SNR frames were initially recorded per movie.

A similar experiment to the above was performed to determine the optimal reference size. As expected, **Figure 4B** confirms that the use of a larger reference leads to a lower *QVal.* From analyzing the *QVal* measurements for different reference sizes and visually inspecting the distortion-vector ERROR fields, we conclude that for our dataset a reference size of 300 × 300 pixels (~3 x ~3 unit-cells) prevented overfitting and allowed beam-induced sample deformations to be accurately corrected.

#### 2.5.2 Combination of different algorithms

Both, the static algorithm Unbend2, and movie-mode algorithms MovieA and MovieB, have advantages for certain image types. If the localized drift is negligible, then Unbend2 has access to the highest SNR for determining the unbending profile, and will give the most reliable results. If frame-internal sample movements are linear, then MovieA is ideal. If instead, such movements are altering direction during dose-fractionated movie recording, then MovieB is the better compromise. A combination of the MovieA and MovieB approaches into one algorithm (linear interpolation at the single frame level of movements determined on super-frames) is possible, but was not yet implemented.

The final algorithms for Unbend2, MovieA and MovieB, including frame averaging, optimal low-pass filtration, and sub-frame-average movie-mode crystal unbending, are illustrated in **Figure 3**.

### 2.6 Merging of low-SNR data

Application of the above algorithms, including image tiling, and three different unbending schemes (Unbend2, MovieA, and MovieB), can lead to a very large number of amplitude-and-phase datasets. If for example a dataset consisting of 400 original movies is to be processed, using 5×5 tiles, then this will yield 400 × 5 × 5 × 3 = 30'000 files that contain amplitude and phase values (so-called APH files), which need to be merged. If each such file contains 1'891 reflections (*e.g.,* if Miller indices range from [0…30; −30…30], and ignoring symmetry-related duplications here), then the merging step will have to merge 56'730'000 reflections, most of which have a minimal SNR, which means that they have large IQ values (Henderson et al., 1990) or abysmally small figure-of-merit (FOM) values. Accurate merging of such data is therefore of importance.

The MRC program *MMBOXA* measures for each reflection in the Fourier transform of the unbent image the amplitude *Amp*, which is measured as the amplitude above the background baseline, the phase *Pha*, and the background *Back* as average of the amplitude values of pixels along the edge of a square around the reflection of interest. Based on the ratio of *Amp* and *Back*, an IQ value is assigned to each reflection (Henderson et al., 1990), as

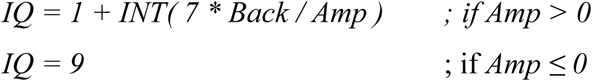

This means that

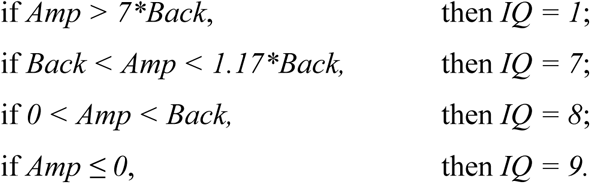

Theses *Amp /Back* ratios can also be translated into a figure-of-merit (FOM) value, as

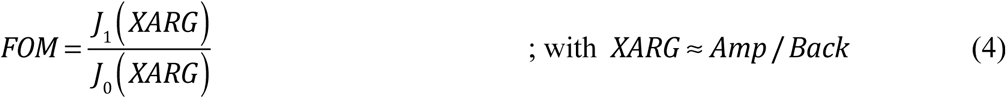

with *J*_1_ and *J*_0_ being the modified Bessel function of the first kind of order 1 respectively 0. The calculation of *XARG* from *Amp* and *Back* is implemented in the MRC program MMBOXA. For details, see (Blow and Crick, 1959; Sim, 1959; Sim, 1960).

The above relation can be inverted computationally through an interpolated look-up table, to give a function *XARG* (*FOM*) = *lookup _ table* (*FOM*)

Amplitude measurements for the same reflections *i* from *n* different APH files can be averaged as

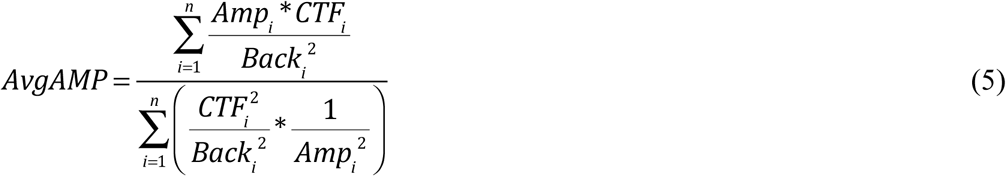

where *Amp* = measured amplitude value for this reflection, *CTF* = fitted CTF value for this reflection, *Back* = measured background amplitude around this reflection.

Phase measurements are averaged as

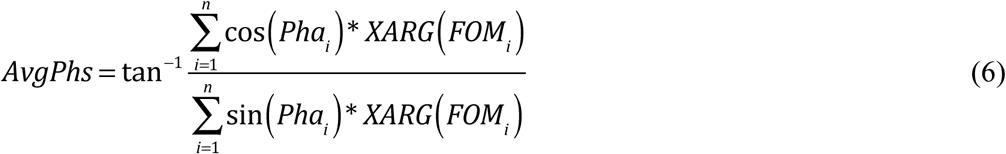

where *Pha* is the measured phase value for this reflection.

The above calculated averages for amplitude and phase values can also approximated by

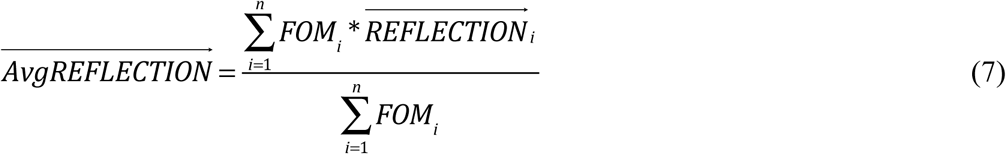

with 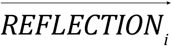 being the complex-valued reflection, consisting of an Amplitude and Phase value (or, alternatively, a *cos* and a *sin* term, which will result in the same averaging). The *FOM* values can be averaged as

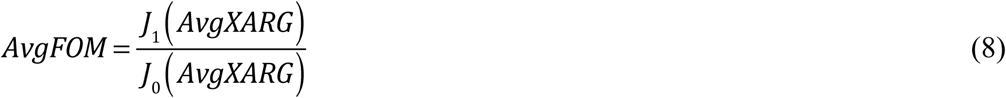

with

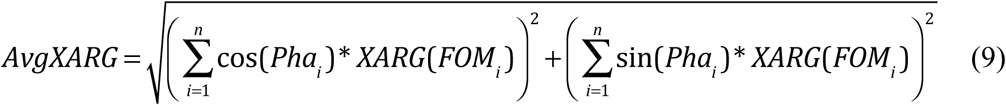

If the phases were all identical, then the average FOM value calculation could be simplified to

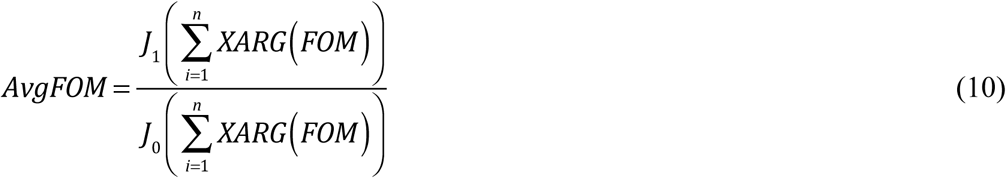

However, in the general experimental case, the measured phases vary, so that (8) and (9) are required.

### 2.7 Merging of reflection data in 3D space into a volume

For a 2D crystal sample of infinite size and perfect order, the unit cell would be infinitely repeated in X and Y directions, but in the vertical Z direction the sample would offer only one single layer of the unit cell density. For this reason, the 3D Fourier space representation of that crystal would have vertically oriented so-called lattice lines, along which amplitude and phase values would smoothly vary, while there would be in the general case no correlation between the values on different lattice lines. This is different from the situation in single particle cryo-EM, where due to the limited extend of the sample not only in Z, but also in X and Y directions, the 3D Fourier space has a neighborhood correlation between adjacent pixels in Fourier space in all directions.

For a 2D crystal, due to the finite support in the Z direction that can be approximated with a rectangle masking function (the crystal has positive density within a certain Z range, but its density is zero above and below that Z range), its Fourier space representation is convoluted in the vertical direction along the lattice lines with a *sinc* function.

A 2D crystal image after unbending will in its Fourier transform show distinct reflections that can be numbered by 2D Miller indices *h,k.* These reflections belong in 3D Fourier space to specific vertical heights *z*^*^ on the lattice lines, whereby the *z*^*^ value is depending on the tilt geometry. A full dataset of 2D crystal images will yield APH files with thousands or millions of such reflection measurements, each belonging to a lattice line *h,k* and a height *z*^*^. Classically, the MRC software used the program *LATLINE* to transform the set of reflections into lattice line data, by fitting a set of *sinc* functions with a least-square error minimization to the experimental values (Agard, 1983).

In our hands, this lattice line *sinc* fitting procedure was difficult to control, when using with a very large dataset of with minimal SNR. We therefore replaced it with a weighted Fourier backprojection, in which the reflections at their *h,k,z^*^* position are inserted into an Euclidian 3D Fourier volume, while computing *FOM*-weighted averages using equation (4) above for Fourier voxels, where several measurements are present. After Fourier inversion, this procedure resulted in a backprojected 3D volume that showed much less noise than when using lattice line fitting. This procedure then invites for computing two half-set 3D reconstructions, which can be compared by Fourier shell correlation (FSC). Different procedures can be implemented in order to attempt reaching semi-independence of the two sub-volumes, such as assigning the even- and odd-numbered APH files to different subvolumes. Similarly to the procedures implemented in RELION, the FSC curve between the two sub-volumes can be used to resolution-limit the 3D reconstruction obtained from a backprojection of the entire dataset, to obtain a 3D volume that automatically has a down-weighted amplitude for frequencies that are less reliable. Such a 3D volume can then be used as alignment reference for the subsequent round of refinement of phase origins for the images during the iterative 3D merging cycles.

### 2.8 Retrieval of missing amplitudes and phases by PCO

Filling the data in the missing cone is a signal recovery problem, where the missing spots in Fourier space are to be recovered using *a priori* information about the object under consideration. Phase retrieval, another signal recovery problem, has attracted a lot of interest (Fienup, 1982; Saldin et al., 2001; Wedberg and Stamnes, 1999; Yan et al., 2011). In this case, the phase of a complex-valued function with known modulus has to be estimated. Many phase retrieval methods rely on Fienup’s approach (Fienup, 1978), where known constraints are applied iteratively in the object space and its Fourier space. Bauschke et. al. (Bauschke et al., 2002) showed that the approaches used by Fienup, namely the *error reduction algorithm*, the *basic input-output* algorithm and the *hybrid input-output* algorithm, can be mathematically modeled as convex optimization problems. This approach was used by Barth et. al. (Barth et al., 1989) to retrieve phases in electron microscopy (EM). In 2D electron crystallography imaging, the problem is slightly different to phase retrieval, since the amplitudes and phases are known for part of the Fourier space, while for the remainder these need to be estimated.

The work by Gipson *et al.,* (Gipson et al., 2011) presents an iterative approach called *projective constraints optimization* (PCO) that addresses this challenge to fill in missing amplitudes and phases. The method relies on iteratively applying known constraints in real and Fourier space. It also provides a transform to convert the uniformly spaced lattice lines in real space to the non-uniform reciprocal space lattice lines, replacing the traditional lattice line fitting (Agard, 1983). Uncoupled amplitudes and phases are used as the raw data, and truncated singular value decomposition is applied to transform between the object space and irregularly sampled Fourier space with fractional steps and individual point-wise weighting.

Here, we extend the convex optimization problem to fill in missing amplitudes and phases in the 3D Fourier space generated from real-space 2D electron crystallography images. Unlike the earlier PCO implementation or traditional lattice like fitting, we employ weighted back-projection to obtain the first reconstruction using density images, and then use this data in an iterative approach to retrieve missing amplitudes and phases. In addition, we introduce the possible constraints in real and reciprocal spaces including a modified version of the shrinkwrap algorithm introduced by Marchesisni *et al.* (Marchesini et al., 2003) for X-ray image reconstruction.

#### 2.8.1 Algorithm

The approach is to develop a sequence of possible solutions by enforcing the known constraints. At a given point in the sequence, the algorithm first fulfills the constraints in the Fourier domain, followed by fulfilling the constraints in the real object domain. This results in a new term of the sequence. The produced term of the sequence is an intersection of the required constraints from Fourier and object space. It should be noted that the terms of the sequence change less as the iterations progress. In other words, the error is reduced and the result converges to a stable solution. Hence, the sequence can be iteratively developed until a convergent solution is found. The proof of convergence of such algorithms can be found elsewhere (Combettes and Trussell, 1990; Il'ina et al., 2009).

*Projection operators* are used to apply the constraints to real and object space. The projection operator that applies object constraints, *P_O_*, can be approximated by setting all elements in the current set *s_n_* that do not lie in the support to zero (Deutsch, 1980). Here, the set *s_n_* refers to the current reconstruction volume of the protein. When applied, the projection operator for Fourier constraints, *P_F_*, will replace the reflections in Fourier space by the set of known reflections. Thus, the expressions to progress the sequence given the current term *s_n_* can be written as

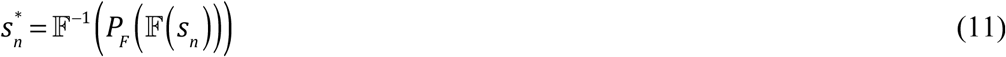

and

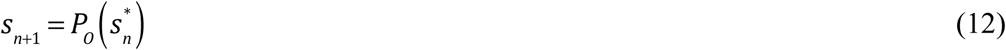

with 𝔽 being the Fourier transform. It should be noted that this scheme is based on Fienup’s *error reduction* approach (Fienup, 1986). Other variants with *basic input output* and *hybrid input output* can also be applied. For example,Deng *et al.* developed a similar approach for electron tomography datasets (Deng et al., 2016). Simulation studies to test the recovery of data in the missing cone region show an excellent performance of this algorithm, at least in the absence of experimental noise **(Supplementary Fig. 3)**.

#### 2.8.2 Extracting the known reflection set

It is essential to have a correct set of reflections to use in the iterative algorithm. As outlined above, in 2D electron crystallography, images of 2D crystals are recorded from the non-tilted and the tilted samples. The image data collected from a series of non-tilted 2D crystals is merged to obtain an initial estimate of the horizontal (X,Y, Z=0) plane of the Fourier transformed object. The image data from tilted crystals are merged onto this plane using the *Central Projection Theorem*, filling values in higher (Z ≠ 0) regions of the Fourier transformed object. For a certain point on a lattice line 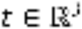, several measurements for Amplitude, Phase, and FOM values might exist, originating from the same or different images, which in that case would have to be combined. This can be achieved using the averaging techniques described above.

#### 2.8.3 Defining the support in object space

An approximation of all the data points (voxels) in object space that are expected to have non-zero values, defines the support of the object. This support set can be developed using the constraints on the density values that a particular object under consideration could achieve. Some of these constraints are demonstrated below.

##### 2.8.3.1 Membrane slab

The target objects of electron crystallography are often 2D crystals of membrane proteins, *i.e.,* sheet-like structures with a repeating unit. This protein sheet has a well-defined boundary in the vertical (Z) axis. One can exploit this fact and restrict the support to a certain length on the Z axis.

##### 2.8.3.2 Real valued, non-negative densities

Protein structures should always be real-valued and non-negative. Consequently, all negative densities can be removed from the support.

##### 2.8.3.3 Modified shrinkwrap optimization

The shrinkwrap optimization updates the support using the density at the current iteration (Marchesini et al., 2003). This algorithm defines a protein envelope by applying a low-pass filter to the current volume, which is subsequently thresholded at a low density to update the support. This assumes that all true protein features can be contained within the blurred low-pass filtered envelope. Densities outside of this envelope are considered to be noise. The threshold is data-set dependent and should be chosen so that the noise is avoided without the loss of true features.

We introduced a modified version of this shrinkwrap optimization, where instead of using a binary mask we use a weighted masking with two different density thresholds ε_1_ and ε_2_, with ε_1_ < ε_2_. Using the above procedure we create a low-pass filtered volume *D_0_*, which is thresholded, forming two sets *D_1_* and *D_2_*, defining the support of the object, where *D_1_* corresponds to the set with threshold ε_1_, and *D_2_*,corresponds to the set with threshold ε_2_. The volume *D_1_* then includes the volume of *D_2_.* With these, the projection operator *P_O_* can be defined as

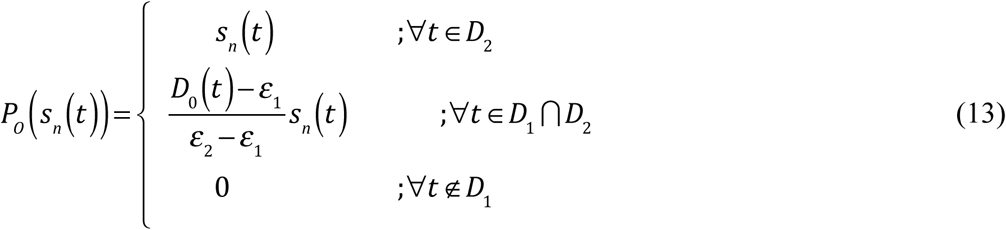

This means that all the densities lying within the inner mask *D_2_* are kept intact, while the densities lying outside the outer mask *D_1_* are set to zero, and the densities between the two masks are linearly scaled down. Using the modified shrinkwrap optimization, one can chose thresholds and minimize the errors that occur with the assumptions of traditional binary masking.

To make this shrinkwrap constraint more conservative, it can be restricted such that it only modifies Fourier voxels within the missing cone region:

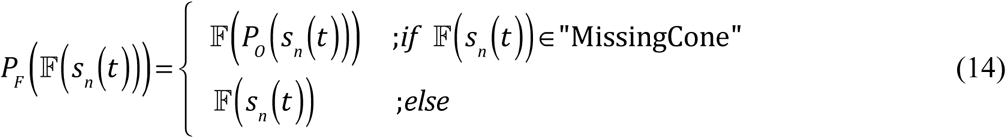

Or, after each iteration, the Fourier voxels outside of the missing cone defined by a given limiting angle are replaced with the original values.

## 3 Results and Discussion

Cryo-EM images of 2D crystals of the cAMP-modulated potassium channel MloK1 (130 × 130 Å unit cell, P42_1_2 symmetry, vitrified on ultrathin carbon film supported by holey carbon film) were used to test the presented algorithms. This four-fold symmetric ion channel undergoes conformational changes upon cyclic adenosine monophosphate (cAMP) binding (Kowal et al., 2014). Here, we used the cryo-EM data recorded in the presence of cAMP (open channel conformation, PDB Model: 4CHV).

### 3.1 Data acquisition and classical image processing

We recorded over 500 dose-fractionated movie-sequences, using a FEI Titan Krios equipped with a Gatan K2 summit detector (without energy filtration), operated at 300kV, and recording dose-fractionated images in super-resolution electron counting mode, yielding movies with 8k × 8k frames. Images of crystals with a nominal sample tilt of up to 50° were recorded. The microscope was operated in low-dose mode at a nominal magnification of 22,500x at the screen level, resulting in an effective magnification of ~37,000x on the detector. The resulting physical pixel size was 1.30 Å on sample level, respectively 0.67 Å for the super-resolution pixels. Movies consisting of 40 frames were recorded over 16 seconds total exposure, resulting in a 0.4 second exposure for each frame. The total electron dose per movie was ~40 electrons/Å^2^, respectively ~1 electron/Å^2^ per frame. Micrograph defocus varied between −0.75 and −4.3 μm. To avoid undercounting (Li et al., 2013a), the electron dose rate was set to 5 electrons per physical pixel per second. The images were corrected for whole-frame-drift with MotionCorr2.1 (Li et al., 2013b) on-the-fly and automatically processed by *2dx_automator* as detailed in (Scherer et al., 2014). The best 346 movies were used for the final 3D reconstruction. These images showed the relatively small 2D crystals typically only within about half the image size. After smooth-edged masking of the crystal areas, the direct Fourier transforms of these images showed the crystal reflections usually upto only 12 Å resolution, in very few cases up to 10Å resolution.

The refinement algorithm described above was applied to the merged 3D dataset. The constraints used in object space included non-negative real densities, a membrane slab and the modified shrinkwrap optimization described above. For the shrinkwrap optimization the volume was first low pass filtered at 10 Å with the higher and lower density thresholds set at respectively 11% and 8% of the maximum density value in the present volume. The data was recovered only in a specific vertical conical region in Fourier voxels belonging to a tilt angle higher than 55 degrees (corresponding to a cone with apex angle of 140 degrees), while the remainder of the Fourier space data was left untouched. In addition, in each iteration the amplitudes of the Fourier voxels in the newly created cone region were scaled so that their average was 75% of the average amplitudes of the Fourier voxels in the higher half of the experimentally sampled tilt angles.

### 3.2 CTF Correction

The effect of the CTF was corrected in stripes before unbending using phase flipping. Alternative correction of the CTF by using TTF correction and other approaches involving CTF correction before and after unbending using Wiener filtration or CTF multiplication were tried, but the best results were obtained using phase flipping in stripes before unbending, as it corrects the phases and at the same time produces least amount of artifacts in the amplitudes.

### 3.3 Structural improvements

The best 346 movies were processed as described above, using Unbend2, MovieA and MovieB routines, followed by 5×5 tiling of the images and recalculation of tilt geometry and defocus values at each tile. A new “final map” was produced from each of these tiles. Due to the limited size of the crystals, the majority of the tile datasets could be deleted, since these tiles did not show any Fourier reflections at all. The remaining tiles with significant signal (a total of 6361 tiles) were considered for the final 3D reconstruction. **Figure 5** shows the quality improvement due to movie-mode unbending for different crystals featuring different tilt angles. The impact of movie-mode unbending on a resulting crystallographic unit cell is shown in **Figure 6**, for both, whole-frame drift-corrected images, using MotionCorr (Li et al., 2013b), and alternatively the newer drift correction MotionCor2 (Zheng et al., 2016).

**Figure 5:**
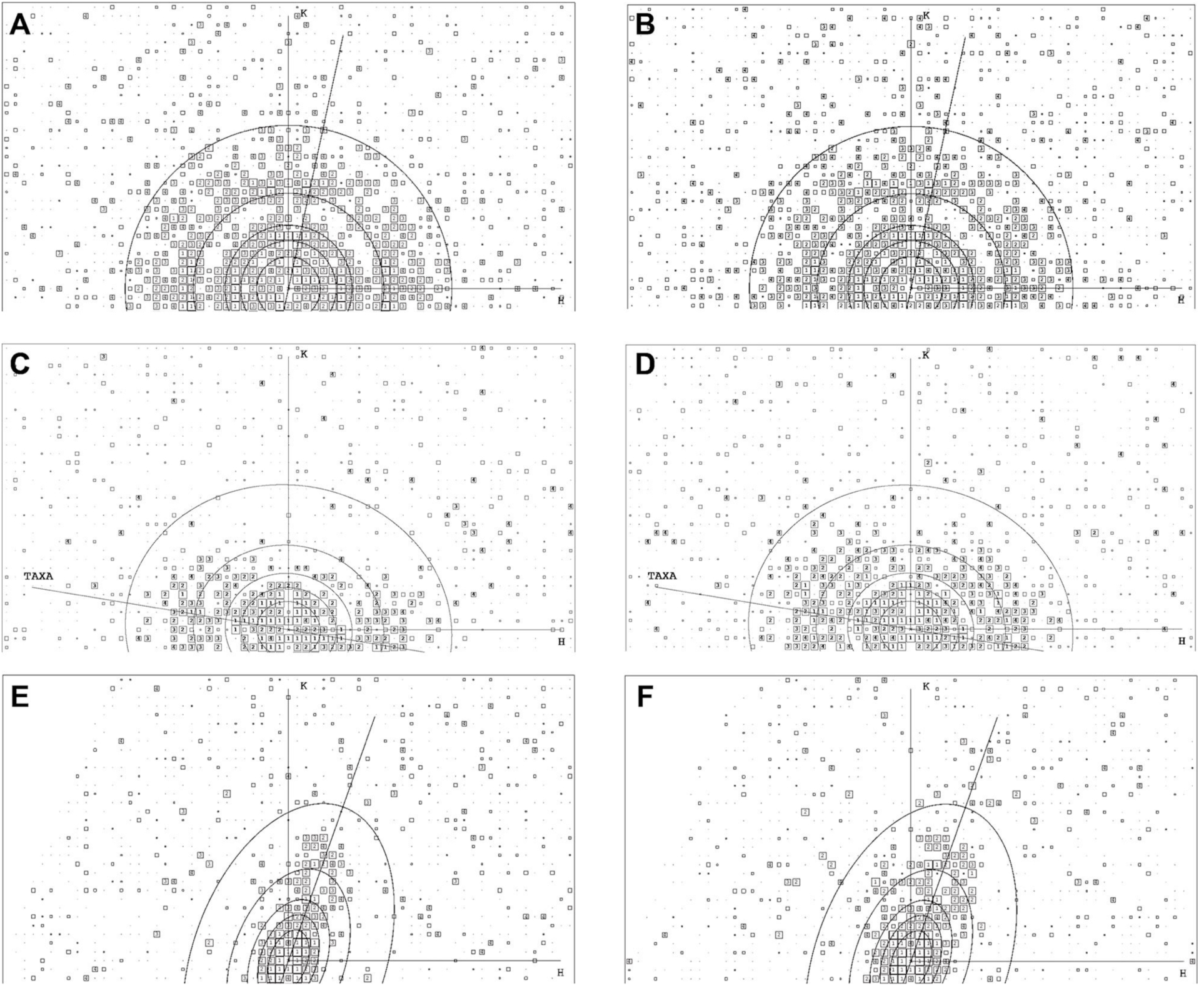
Canonical IQ-plots obtained without and with movie-mode unbending applied to crystals with different tilt angles. The Nyquist frequency of these plots is at 4 Å; the resolution circles are at 36 Å, 24 Å, 18 Å, 12 Å and 7 Å. (**A**) IQ-plot of a non-tilted 2D crystal without movie-mode unbending, *i.e.,* unbending with Unbend2 of the drift-corrected sum of all movie frames. (**B**) IQ-plot of the same exposure as in (A) but after movie-mode unbending with MovieB. (**C**) IQ-plot of a 20° tilted crystal without movie-mode unbending and (**D**) IQ-plot after unbending with MovieB. IQ-plot of a 50° tilted crystal (**E**) without movie-mode processing and (**F**) after unbending with MovieB. It can be seen that the IQ-plots after movie-mode processing show significantly more spots and also spots at higher resolution, especially orthogonally to the tilt axis (labeled with “TAXA”).

**Figure 6:**
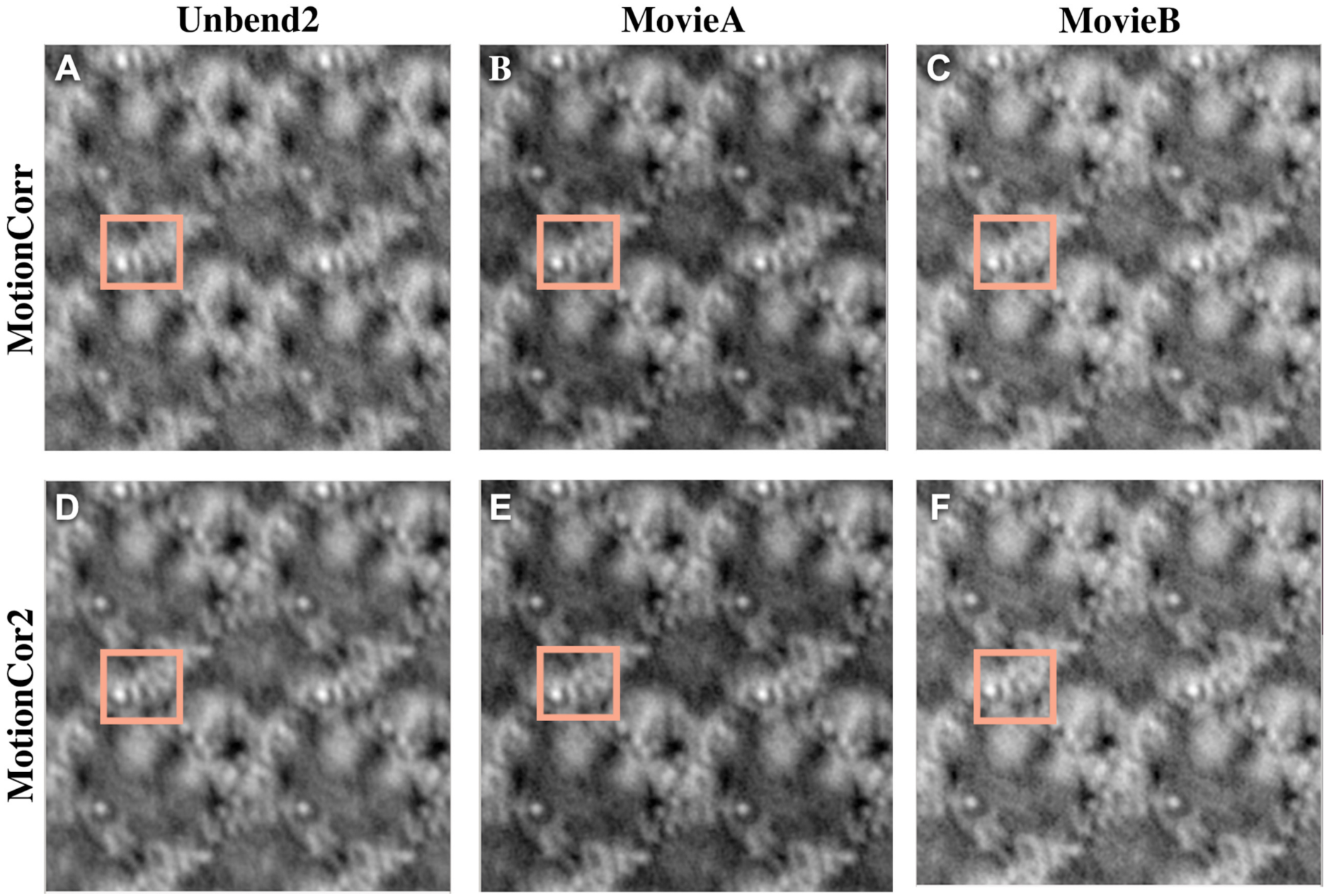
Movie-mode unbending applied to an image of a 45° tilted crystal compared to classical processing using different drift correction software MotionCorr and MotionCor2. (**A**)Projection map obtained by drift correction with the whole-frame drift correction software MotionCorr (Version 2.1, (Li et al., 2013b)) and classical processing using Unbend2. (**B**) Projection map produced by drift correction with MotionCorr and movie-mode unbending using the MovieA algorithm. (**C**) Projection map produced by drift correction with MotionCorr and movie-mode unbending using the MovieB algorithm. (**D**) Projection map obtained by drift correction with the image-warping capable drift correction software MotionCor2 and classical processing using Unbend2. (**E**) Projection map produced by drift correction with MotionCor2 (using the option *-Patch 5 5,* (Zheng et al., 2016)) and movie-mode unbending using the MovieA algorithm. (**F**) Projection map produced by drift correction with MotionCor2 and movie-mode unbending using the MovieB algorithm. The tilt axis is roughly horizontal. Inferred from the densities visible, the movie-mode processing resulted in higher resolution than the classical unbending. For instance, two helices in the marked square are separated in (B) and (C), but not in (A). MotionCor2 produces better results than MotionCorr and the movie-mode unbending appears to marginally improve the visibility overall sharpness of the image. The unit cells of these 2D crystals in p42_1_2 symmetry have dimensions of 135 × 135Å with a 90° angle. These panels show 2×2 unit cells, under 45° sample tilt.

Merging of the data from whole-frame drift-corrected movies using MotionCorr, and then applying Unbend2, MovieA and MovieB, and refinement of tilt geometries on tiles, iterative refinement of the final volume resulted in a 3D map reproduced in **Figure 7**. The final map resolves all alpha-helical elements of the potassium channel, including the voltage sensor domains S1-S4, the channel helices S5 and S6, and the horizontal C-linker helix. The C-terminal cyclic nucleotide binding domain (CNBD) contains alpha-helical elements and also a horizontally oriented beta-sheet clamp, which is partly resolved (**Figure 7C**). This structure is detailed elsewhere (Kowal et al., 2017). This map shows significant improvement over an earlier map, produced from the same dataset but without the here described algorithmic improvements (**Figure 7D,E**, (Scherer et al., 2014)), and is strongly improved over earlier film data (**Figure 7F,G**, (Kowal et al., 2014)).

**Figure 7:**
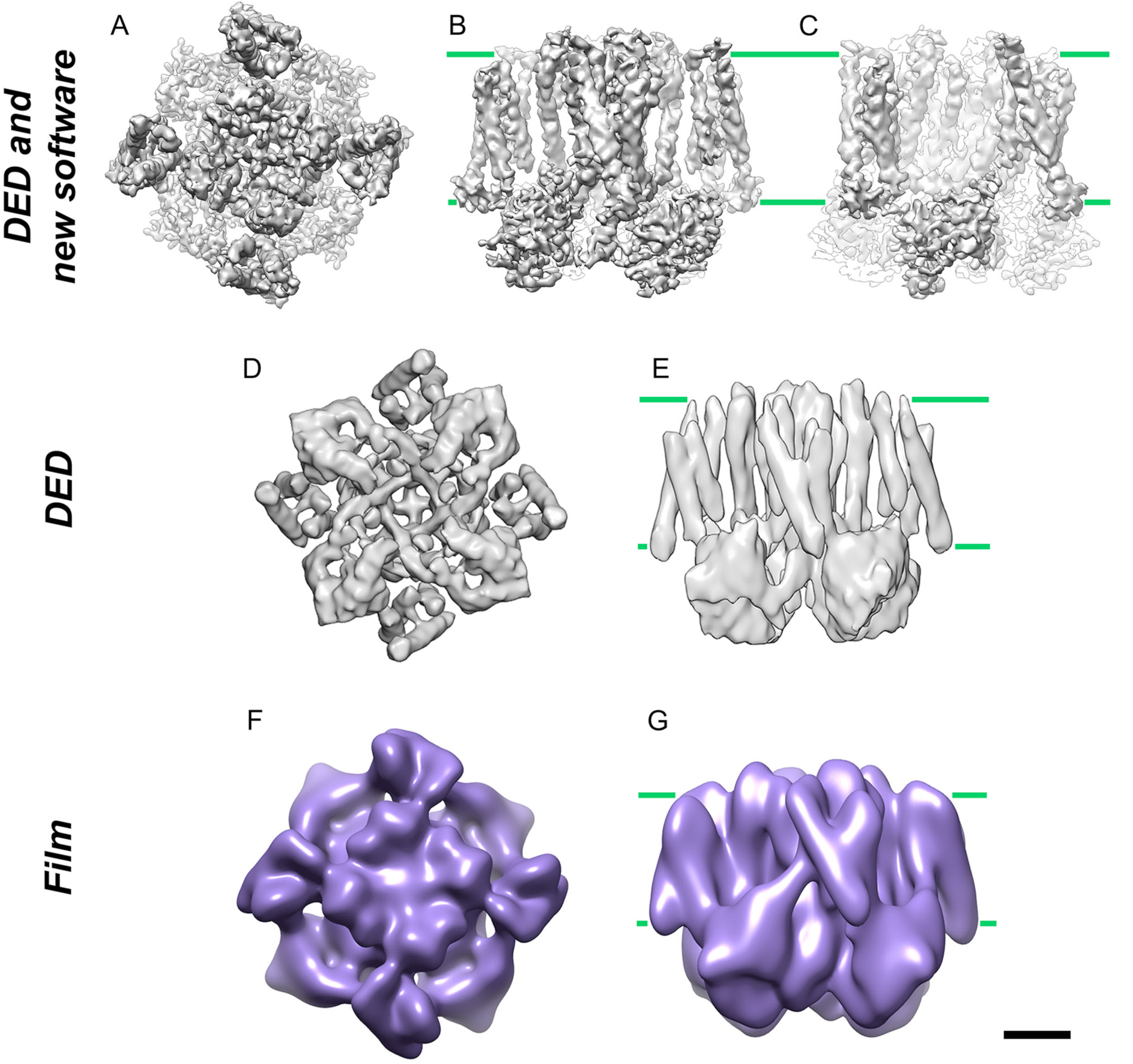
Comparison of MloK1 3D-density maps. (**A**) Top views of the 3D map generated using 2D crystal images obtained from FEI Titan Krios equipped with a Gatan K2 summit direct electron detector (without energy filtration) and subsequent application of the algorithms stated in this work. (**B**) Side-view of the reconstruction in A. The green lines indicate the approximate boundaries of the membrane plane. (**C**) same as (B), rotated by 45° in the membrane plane. (**D**) Top view and (**E**) side view of the 3D map from previous work on the same 2D crystals with the same microscope and K2 Summit camera, but before development of the here described algorithms (Scherer et al., 2014). (**F**) Top view and (**G**) side view of the 3D map from previous work on the same 2D crystals with the same microscope, but obtained using photographic film (Kowal et al., 2014). Scale bar for all panels 20 Å.

### 3.4 Algorithmic alternatives

Averaging batches of sequential movie frames into super-frames to reduce the risk of overfitting is the key point of the MovieB algorithm. Its drawback is that processing these super-frames produced distortion-vector ERROR fields that are a compromise for the averaged frames. The distortion-vector ERROR plots calculated for different super-frames differed significantly from one to the next, so that they could not be interpreted by smooth underlying physical movements. An alternative to the applied grouping into super-frames would be to compute so-called “rolling average” (RA) super-frames, where each original frame would contribute to multiple RA-super-frames. The average built for RA-super-frame (n) would then also include down-weighted contributions from frames (n-2), (n-1) and frames (n+1), (n+2), etc., which could be used with a Gaussian weighting scheme centered on frame (n). The determined ERROR field for each RA-super-frame could then be applied to the single central frame only. Preliminary experiments with this method applied to the presented test dataset did, however, not yield better results than the algorithm described before, possibly because the rolling average images might not have been sufficiently dominated by the central frame to produce a valid ERROR field for that frame. At the extremely low SNR of the individual frames recorded here, computing ERROR fields that are valid for just one frame failed.

## 4 Conclusions

The described algorithms represent procedures tailored to achieve high resolution for cryo-EM images of 2D crystals that show only limited crystalline order. Tiled image processing enables to improve the tilt geometries over the image, if the crystal distortions are severe, or if the 2D crystals cannot be prepared as perfectly flat planes. Movie-mode unbending offers an efficient and generic approach to correct for beam-induced translational motion within the dose-fractionated image frames, which previously limited the quality of 3D reconstruction obtained. We reduced the risk of overfitting by either constricting unit cell movements to linear trajectories only in the MovieA algorithm, or by averaging multiple drift-corrected movie frames into super-frames in the MovieB algorithm. We also employed dose-dependent resolution filters in forms of B-factors to allow processing data with higher electron doses. For the MovieB algorithm, we investigated the properties of its two key parameters, namely the “number of frames to average” and the “reference size”. We also present an easy to implement iterative algorithm to generate the missing data in Fourier space. The algorithm relies on constraints enforced in orthogonal object and reciprocal space, which are iteratively applied to an experimental set of reflections. A volume that satisfies all of the constraints simultaneously is a possible solution of the system. Application of the new software to a real-world dataset significantly improved the resolution of the final 3D map, so that from 2D crystals diffracting to only 10 Å at best, the final 3D reconstruction was able to resolve individual beta strands in horizontally oriented beta sheets, suggesting that a vertical resolution of better than 5.4 Å had been reached.

## 5 Acknowledgments

We thank Shirley A. Müller for insightful discussions and critically reading of the manuscript, and Marcel Arheit for fruitful discussions in the early project stage. This work was supported by the Swiss National Science Foundation (grants 315230_146929, 205320_144427, and the NCCR TransCure).

All the here-described procedures are implemented in the Focus software package, which is available under the Gnu Public License on http://www.focus-em.org.

